# Transcriptional coactivator MED15 is required to maintain β-cell maturity

**DOI:** 10.64898/2026.08.26.745805

**Authors:** Rachel J. Spencer, Meixia Dan, Sara P. Cristiano, C. Bruce Verchere, Francis C. Lynn, Stefan Taubert

**Affiliations:** Centre for Molecular Medicine and Therapeutics, University of British Columbia, Vancouver, BC, Canada; Department of Medical Genetics, University of British Columbia, Vancouver, BC, Canada; Edwin S.H. Leong Centre for Healthy Aging, University of British Columbia, Vancouver, BC, Canada; Diabetes Research Program, Childhood Diseases Research Theme, BC Children’s Hospital Research Institute, Vancouver, BC, Canada; Department of Surgery, University of British Columbia, Vancouver, BC, Canada; Department of Pathology and Laboratory Medicine, University of British Columbia, Vancouver, BC, Canada; School of Biomedical Engineering, University of British Columbia, Vancouver, BC, Canada

**Keywords:** Mediator complex, diabetes, β-cell, maturation, Pax6, Nkx6-1, Med15

## Abstract

The Mediator complex, a vital transcriptional coregulator in eukaryotes, partners with transcription factors to orchestrate gene transcription and in turn many developmental and physiological processes. Mediator subunit MED15 is required for the pre-natal development and post-natal maturation of pancreatic β-cells in mice. However, whether MED15 plays a role in β-cell function after initial development and throughout adulthood is unknown. To investigate the role of MED15 in β-cells post-maturation, we induced a β-cell specific *Med15* knockout at six weeks age in male and female mice. This post-developmental *Med15* ablation led to glucose intolerance and impaired insulin secretion. RNA-sequencing revealed downregulation of β-cell maturation markers, indicating that MED15 is continuously required to maintain β-cell functionality. Further, we implanted insulin pellets into *Med15* knockout mice to lower blood glucose and used RNA-seq to validate that the transcriptional changes we observed are a direct consequence of *Med15* loss and not an indirect effect of hyperglycemia arising in the knockout mice. In sum, our study shows that MED15 is continuously required after weaning to maintain functional β-cell maturity.

**Article Highlights:**

- Mediator complex subunit MED15 is required for post-natal β-cell maturation, but its role in adult β-cells was unknown
- Ablation of *Med15* in β-cells of adult mice resulted in glucose intolerance and loss of maturation
- β-cell maturity and transcriptional defects are not rescued by controlling glycemia with insulin implants
- MED15 is required to maintain β-cell maturation post-weaning and sustain β-cell function throughout life

## Introduction

In humans and in some animals, autoimmune destruction of pancreatic β-cells and loss of insulin secretion lead to type 1 diabetes, whereas β-cell dysfunction combined with peripheral insulin resistance causes type 2 diabetes. At birth, β-cells are differentiated but remain immature, defined as having a low threshold for glucose-stimulated insulin secretion (GSIS) (1). Immature β-cells then undergo a maturation phase between birth and weaning to become fully functional and glucose responsive (2). β-cells must then maintain functional maturity throughout life, otherwise hyperglycemia and glucose intolerance ensue (3).

Maturation is driven by a genetic network featuring transcription factors. Some, like Pancreatic and Duodenal Homeobox 1 (PDX1), Neurogenic Differentiation 1 (NEUROD1), and NK6 homeobox 1 (NKX6-1) are required from the early stages of pancreas development to maintaining maturation post-weaning (4–6). Others, like v-maf musculoaponeurotic fibrosarcoma oncogene homolog A (MAFA), are upregulated during and involved in driving maturation, and loss of their expression causes loss of maturation (7). Finally, some proteins are dispensable for initial maturation, but are expressed afterwards in mature β-cells and serve as maturity markers, e.g. Urocortin-3 (UCN3) (8). The importance of the transcriptional circuits that drive β-cell development, differentiation, and maturation is highlighted by the fact that mutations in some transcription factors cause early-onset, monogenic forms of diabetes (9).

Transcription factors require transcriptional coregulators to regulate gene expression (10). One critical coregulator is the Mediator complex, which is conserved across eukaryotes (10). Mammalian Mediator is composed of ∼30 subunits and adopts a conserved architecture featuring the Head, Middle, Tail, and Kinase modules (11,12). Functionally, Mediator enables RNA polymerase driven gene transcription by acting as a molecular bridge between transcription factors and polymerases, by regulating polymerase pausing, elongation, and termination, and by forming chromatin loops (13).

Interestingly, Mediator subunits control metabolism in several species (14). MDT-15/MED15 coordinates lipid metabolism by interacting with sterol regulatory element-binding proteins (SREBP) in Caenorhabditis and in mammals (15–17). In mice, MED23 regulates gluconeogenesis through Forkhead box protein O1 (FOXO1) in the liver (18) and MED13 inhibits glucose uptake in skeletal muscle (19). Cyclin-Dependent Kinase 8 (CDK8) promotes glucose uptake and glycolysis in cancer cells (20) but negatively regulates β-cell insulin secretion by phosphorylating OSBPL3 (21). Finally, in mice, MED15 is required for post-natal β-cell maturation by interacting with NKX6-1 and NEUROD1 and promoting the expression of genes involved in maturation (22). Accordingly, mice with β-cell specific *Med15* loss at birth have impaired glucose uptake, insulin granule maturation, insulin secretion, and glucose tolerance. Thus, MED15 is required within β-cells to attain maturation and function between birth and weaning.

Because *Med15* is expressed throughout life, and because islets from humans with T2D have reduced *Med15* expression (22), we hypothesized that MED15 might be continuously required in β-cells even after maturation. To study MED15 in adult β-cells, we generated an inducible knockout mouse model and explored the phenotypic and transcriptomic consequences of *Med15* ablation after weaning. *Med15* knockout mice develop glucose intolerance due to dysfunctional insulin secretion and loss of β-cell maturity. Our results show that MED15 is required throughout life to maintain β-cell function and maturity.

## Research Design and Methods

### Animal studies

All mouse experiments were approved by the University of British Columbia Animal Care Committee. *Med15^fl/fl^* (RRID:MMRRC_058472-UCD), *mTmG^fl/fl^* (RRID:IMSR_JAX:007576), and *Pdx1^CreER/+^* (RRID:IMSR_JAX:024968) mouse lines were used, and maintained on a C57BL/6J background (22–24). Control mice were *Med15^fl/fl^; mTmG^fl/fl^; ^+/+^* and experimental mice (M15KO) *Med15^fl/fl^; mTmG^fl/fl^; Pdx1^CreER/+^*. Animals were housed under a 12-h light/dark cycle, fed ad libitum with a standard chow diet (5010; Lab Diets).

At 6 weeks of age, mice received 8mg of tamoxifen (TRC-T006000; Toronto Research Chemicals) dissolved in corn oil via oral gavage every other day for 5 days. Unless otherwise stated, pancreas or islets were collected for analysis two weeks after the final tamoxifen dose.

Mice in the insulin pellet or sham treatment cohort received Meloxicam (5mg/kg) before anesthetization with isoflurane (2% mix with 2L/min oxygen flow). The skin was pierced with a 16G disposable hypodermic needle, then a sterile trocar (12G) containing the insulin implant (As-1-L; LinBit; LinShin, Toronto, Canada) was inserted. For sham treatment, the same procedure was followed except an empty trocar was inserted.

### Islet isolation

Mice were euthanized with isoflurane followed by cervical dislocation. Islets were isolated through collagenase digestion and then left to recover overnight in RPMI1640 (11875093; Gibco) supplemented with 10% FBS (12483020; Gibco), 1% penicillin streptomycin (15140-122; Gibco) and 2mM L-glutamine (35050-061; Gibco).

### Intraperitoneal glucose tolerance test

Following a 12-h fast during the dark cycle, mice were weighed and fasting saphenous vein blood glucose levels were obtained using a OneTouch UltraMini glucometer (Lifescan). Before knockout and two and six weeks after knockout, 2g/kg of 20% w/v D-glucose were delivered via intraperitoneal injection. At ten weeks after knockout 1g/kg D-glucose was used. Blood glucose levels were measured at 10, 30, 45, 60, and 90 min after injection. Blood was collected at fasting and 15min after injection for determination of plasma insulin levels using ELISA. Blood glucose levels greater than the glucometer detection limit are reported as 33.3mM.

### Islet perifusion

Perifusion of isolated islets was performed using a Perifusion V2.0.0 system (Biorep). Islets were perifused with 2.8mM glucose in Krebs-ringer bicarbonate HEPES (KRBH; 114mM NaCl, 20mM HEPES, 4.7mM KCl, 2.5mM CaCl_2_, 1.2mM KH_2_PO_4_, 1.17mM MgSO_4_, 0.2% BSA, pH7.4), followed by 16.7mM glucose, then 30mM KCl plus 2.8mM glucose. Females had an additional perifusion step with 10mM α-ketoglutarate plus 2.8mM glucose before KCl. Following perifusion, islets were transferred to 500μL of acid ethanol (0.1M HCl in 70% ethanol in H_2_O) and stored overnight at 4°C. The cell pellet was used for DNA extraction using Qiagen DNEasy Blood and Tissue kit (69504; Qiagen) and DNA concentration measured on Nanodrop (ND-1000).

### Total insulin content

After overnight recovery, 10 size-matched islets per mouse were added to 500μL of acid ethanol (0.1M HCl in 70% ethanol in H_2_O) and stored overnight at 4°C. Following centrifugation, the supernatant was used to measure total insulin using ELISA.

### Insulin ELISA

A chemiluminescent ELISA kit (80-INSMR-CH01; ALPCO) was used according to the manufacturer’s instructions. Serum insulin samples and perifusion samples were used directly; acid ethanol total insulin samples were diluted 50x. Briefly, 5μL per well of samples and standards were loaded, with standards for each separate plate used. Samples were incubated for 2 hours on an orbital shaker at 850RPM with insulin-antibody conjugate, washed 6 times, and incubated with substrate for 1 min. ELISA plates were read on a POLARstar Omega microplate reader (BMG Labtech) with 1s integration time/well.

### RNA isolation and RT-qPCR

RNA isolation and RT-qPCR were performed largely as described (22), see supplementary methods.

### RNA sequencing

RNA sequencing and analysis was performed largely as described (25), see supplementary methods. RNA-seq data are available at Gene Expression Omnibus (https://www.ncbi.nlm.nih.gov/geo/) record GSE343956. Functional enrichment analysis and visualization were performed with easyGSEA and easyVizR in the eVITTA toolbox (https://tau.cmmt.ubc.ca/eVITTA/) (26).

To reanalyze published *Med15*KO RNA-seq data (22), raw reads were downloaded from Sequence Read Archive (SRA) accession PRJNA564631, extracted using fastq-dump, processed and analyzed as above. Venn diagrams of gene set overlaps and associated p-value were generated as described (27). Additionally, published RNA-seq datasets from (6,28–31) were analyzed with eVITTA (26) when available, otherwise processed as above.

### Immunostaining

Immunostaining was performed on paraformaldehyde-fixed, paraffin embedded tissues as described (32), see supplementary methods for detail.

### Quantification and statistical analysis

Statistical analyses were performed with Prism 9.0 (GraphPad Software). Statistical significance was determined using parametric (Student’s t-test or ANOVA) tests with appropriate post-hoc tests, see figure legends. A p-value < 0.05 was considered as statistically significant difference between groups. Non-significant changes are not shown. *p <0.05, **p ≤0.01, ***p ≤0.001, ****p ≤0.0001. Individual P values area available in Supplementary Table S1. Data are presented as mean ± SD.

### Data and Resource Availability

All data generated or analyzed during this study are included in the published article and its online supplementary files and GSE343956.

## Results

### *Med15* deletion in adult mice causes impaired glucose tolerance and insulin secretion

MED15 is required for initial β-cell development and post-natal maturation (22), but whether it is required to sustain β-cell maturity and/or function throughout life is not known. To examine MED15’s function in adult β-cells, we created an inducible, β-cell specific *Med15* knockout mouse model, *Med15 ^fl/fl^; mTmG ^fl/fl^; Pdx1Cre^ER^/^+^*(M15KO) vs. *Med15 ^fl/fl^; mTmG ^fl/fl^; +/+* controls. Knockout induction at 6 weeks of age with tamoxifen (Fig. 1A) resulted in reduced *Med15* at mRNA and protein levels, as expected (Supplementary Fig. 1).

**Figure 1:**
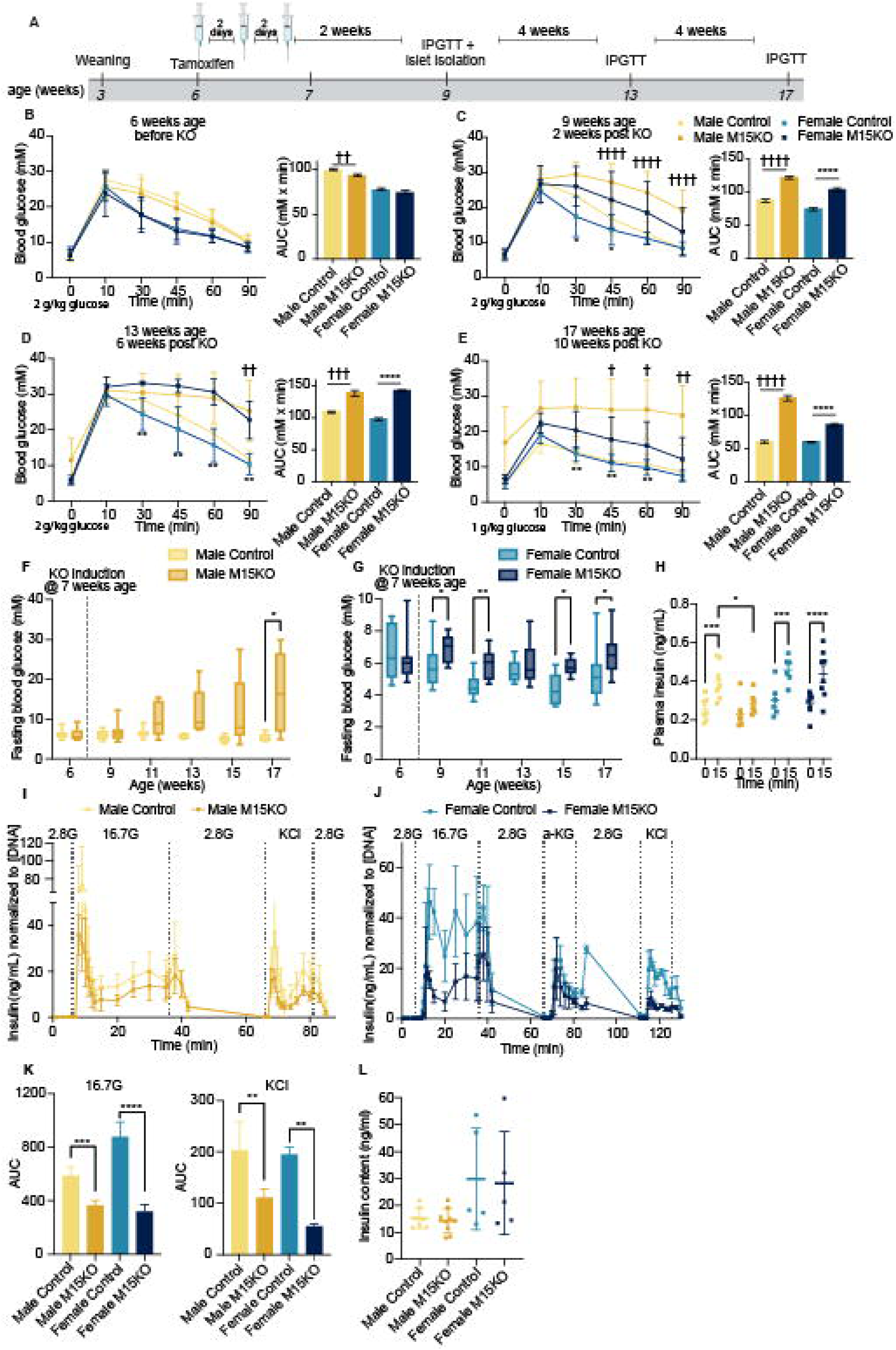
*Med15* knockout in β-cells of adult mice impairs glucose homeostasis. (A) Timeline of Med15 knockout induction. (B-E) Intraperitoneal glucose tolerance test performed after a 12 hour fast and corresponding area under the curve (AUC) (B) before knockout induction (female M15KO n=15, female control n=13, male M15KO n=15, male control n=15), (C) two weeks after KO induction (female M15KO n=12, female control n=10, male M15KO n=12, male control n=8), (D) 6 weeks after KO induction (female M15KO n=10, female control n=9, male M15KO n=5, male control n=5), and (E) 10 weeks after KO induction (female M15KO n=15, female control n=12, male M15KO n=10, male control n=5). 2g/kg glucose were injected in (B,C, and D); 1g/kg glucose was injected in (E). (F-G) Blood glucose levels in males (F) and (G) females measured after 12 hour overnight fast. (H) Serum insulin levels after fasting and 15min after i.p. glucose injection (female M15KO n=8, female control n=6, male M15KO n=7, male control n=7). (I,J) Insulin secretion from isolated islets from (I) male and (J) female mice subject to perifusion of 2.8mM and 16.7mM glucose, 30mM KCl, and 10mM α-ketoglutarate (n=4). The scale of the segmented y-axis in (I) is 70% (0-50) and 30% (65-120). (K) AUC for (I) and (J) for high glucose and KCl perifusion. (L) Total insulin content from 10 islets (female M15KO n=5, female control n=5, male M15KO n=9, male control n=7). In all panels, data are mean ± SD and compared using unpaired Student’s t-test or 2-way ANOVA corrected for multiple comparisons using the Tukey method. Only comparisons between control and M15KO within each sex are shown. * indicates significance in female mice, † indicates significance in male mice.

To test if *Med15* loss compromised glucose tolerance, we performed intraperitoneal glucose tolerance tests (IPGTT). Prior to tamoxifen administration, IPGTTs showed no difference in glucose tolerance between genotypes (Fig. 1B). Two weeks after knockout induction, both male and female M15KO mice were glucose intolerant compared to control (Fig. 1C). Glucose intolerance in M15KO mice of both sexes was sustained until the final time point (Fig. 1D and E). Fasting blood glucose levels were significantly elevated at 10 weeks after knockout in males (Fig. 1F) and at 2, 4, 8, and 10 weeks after knockout in females (Fig. 1G). Male M15KO mice also had decreased plasma insulin levels after glucose injection (Fig. 1H). Male glucose intolerance was more severe than females, potentially reflecting metabolic differences of the B6J background (33). Altogether, M15KO mice have impaired glucose tolerance compared to their control littermates.

The glucose intolerance in M15KO mice could reflect cellular or systemic defects. To test if insulin secretion was intrinsically affected in β-cells by *Med15* loss, we isolated islets from control and M15KO mice and performed *in vitro* perifusion assays. These experiments revealed that M15KO islets secrete less insulin both in response to 16.7mM glucose and when stimulated with KCl, which directly depolarizes the cell membrane (Fig. 1I-K). However, there was no difference in total insulin content between control and M15KO islets (Fig. 1L). These data show that *Med15* is required for pancreatic β-cells to perform insulin secretion.

Glucose intolerance could also be caused by loss of β-cells (34). To assess this in the M15KO mice, we quantified β- and α-cell counts in control and M15KO pancreata. There was no significant difference in the number of INS- or GCG-expressing cells per total pancreatic nuclei, suggesting that β-cell proportion is not affected by loss of *Med15* (Fig. 2A and B). Islet architecture was also unchanged (Supplementary Fig. 2). Thus, the glucose intolerance phenotype in β-cells lacking *Med15* likely arises from functional defects, such as defective glucose sensing, or impaired insulin processing or secretion.

**Figure 2:**
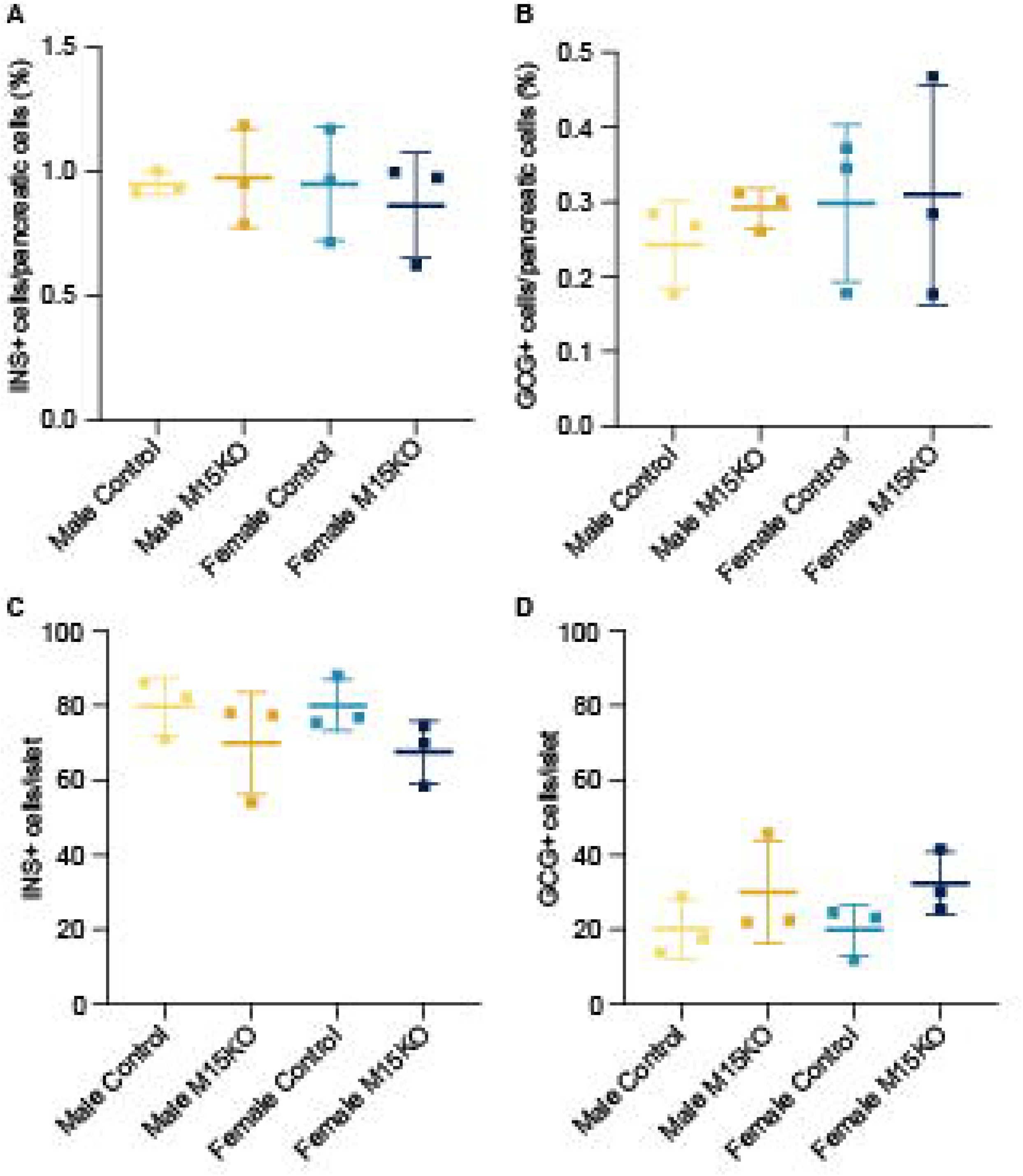
β-cell area is unchanged after *Med15* loss. Quantification of (A) insulin-positive and (B) glucagon-positive cells per DAPI-positive pancreatic cells in male and female M15KO and control mice. Quantification of (C) insulin-positive and (D) glucagon-positive cells per islet. N=3 mice, 4-6 sections per mouse (>250μm intervals). Data are mean ± SD and compared using unpaired t-test.

### β-cell maturity is lost after *Med15* ablation

Because MED15 is a transcriptional coregulator, we next investigated which genes it regulates in the β-cell by performing RNA-seq on whole islets from M15KO and control mice two weeks after knockout (Fig. 3A-B, Supplementary Fig. 3A). Gene expression comparison between M15KO and control islets revealed differential expression of 619 genes in males (279 downregulated and 340 upregulated genes; FDR<0.05, |logFC|≥0.5; Supplementary Table 2), whereas in female M15KO islets, 682 genes were differentially expressed (259 downregulated and 423 upregulated genes; (FDR<0.05, |logFC|≥0.5), Supplementary Table 3), with 117 genes downregulated and 135 genes upregulated in both sexes (Supplementary Fig. 3B). Notably, expression of Mediator subunits other than *Med15* was largely unchanged (Supplementary Fig. 4).

**Figure 3:**
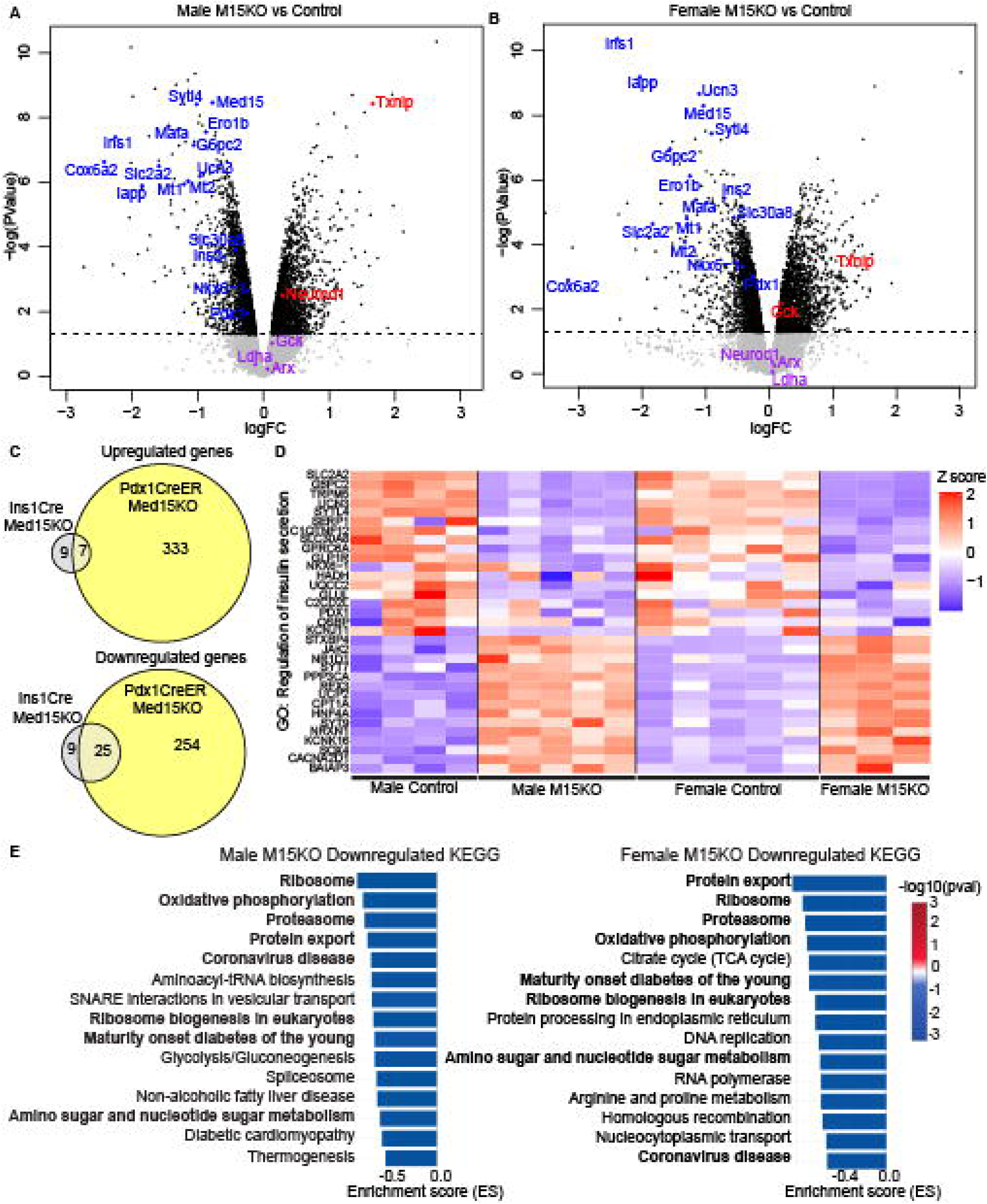
*Med15* is required to maintain the expression of genes involved in β-cell maturity. (A,B) Volcano plots of differentially expressed genes from RNA-seq of whole islets after *Med15* KO in (A) male (control n=4, M15KO n=5) and (B) female mice (control n=3, M15KO n=5) (blue, upregulated; red, downregulated). (C) Venn diagrams showing the overlap of genes regulated by MED15 in post-natal (*Ins1Cre*, grey) and adult (*Pdx1CreER*, yellow) KO models. Numbers indicate the number of differentially expressed genes (FDR < 0.05, logFC < −0.5 or >0.5). (D) Heatmap of dysregulated genes from the gene ontology (GO) pathway ‘regulation of insulin secretion’. Blue indicates decreased gene expression and red indicates increased expression. (E) Top 15 downregulated KEGG pathways in M15KO male and female mice. Bolded pathways appear in both sexes.

To assess if *Med15* deletion in adult β-cells affects gene expression similarly to ablation before birth, we compared our RNA-seq profiles to gene expression profiles seen in our *Ins1-Cre Med15*KO model (GSE137145; male mice only) (22). We reanalyzed the *Ins1-Cre Med15*KO RNA-seq data with our current pipeline to minimize errors due to data analysis and processing. We observed a much higher number of differentially regulated genes (FDR < 0.05) in adult M15KO model (340 up, 279 down) compared to the *Ins1-Cre Med15*KO model (16 up, 36 down) (Fig. 3C). Importantly, however, many β-cell maturation factors and identity genes were downregulated in both *Med15* knockout models (Supplementary Table 4), highlighting the important role MED15 plays in both establishing β-cell maturation and in maintaining the expression of genes after weaning.

The top downregulated genes in both sexes of M15KO included well-known β-cell maturation factors, including *Iapp*, *Ucn3*, *Slc2a2* (which encodes glucose uptake transporter GLUT2), and *Mafa* (Fig. 3A-B). These genes are required to maintain β-cell maturation (7) and/or expressed exclusively in mature β-cells (1). We also observed a dysregulation of numerous genes in the Gene Ontology term “regulation of insulin secretion” in both male and female M15KO (Fig. 3D), in line with the glucose intolerance and impaired insulin secretion seen in M15KO mice. In sum, MED15 is required to maintain expression of mature β-cell transcriptional networks in adult mice.

To delineate pathways affected by *Med15* loss, we performed Gene Set Enrichment Analysis (GSEA). The 15 most significantly downregulated KEGG pathways in both males and females included ‘maturity onset diabetes of the young’ and metabolic pathways such as ‘oxidative phosphorylation’ and ‘glycolysis’ (Fig. 3E). Indeed, *Cox6a2*, a subunit in the electron transport chain, is one of the most strongly downregulated genes in both male and female M15KO islets (Fig. 3A-B). GSEA analysis also identified ‘protein export’ among top downregulated KEGG pathways in both sexes (Fig. 3E), and ‘protein processing in endoplasmic reticulum’ in females, so we further examined endoplasmic reticulum related processes and pathways. Notably, the Biological Pathway terms ‘ER-UPR’, ‘response to unfolded protein’, and ‘response to ER stress’ were downregulated after *Med15* loss (Supplementary Table 6), and genes in the ER-UPR *Eif2a* branch, but not other branches, were also downregulated (Supplementary Table 7). Finally, because loss of β-cell maturity can be caused by interference with the mitochondrial integrated stress response (35) we evaluated these ISR genes, but they are largely downregulated or unchanged (Supplementary Fig. 5). Thus, the loss of maturity in M15KO mice is likely not an indirect consequence of ISR disruption.

We validated the downregulation of several maturation genes and proteins with RT-qPCR and immunofluorescence, confirming that Slc2a2 and Ucn3 have decreased expression at mRNA and IAPP and UCN3 at protein level in the M15KO islets (Fig. 4). Furthermore, we compared our RNA-seq dataset to a published cluster of immature β-cell genes generated from single-cell RNA-seq (36), which revealed increased expression of these immaturity genes in M15KO islets (Supplementary Fig. 6). This supports the notion that M15KO islets have an immature-like transcriptome. Overall, these data demonstrate that β-cells lose maturity after *Med15* deletion.

**Figure 4:**
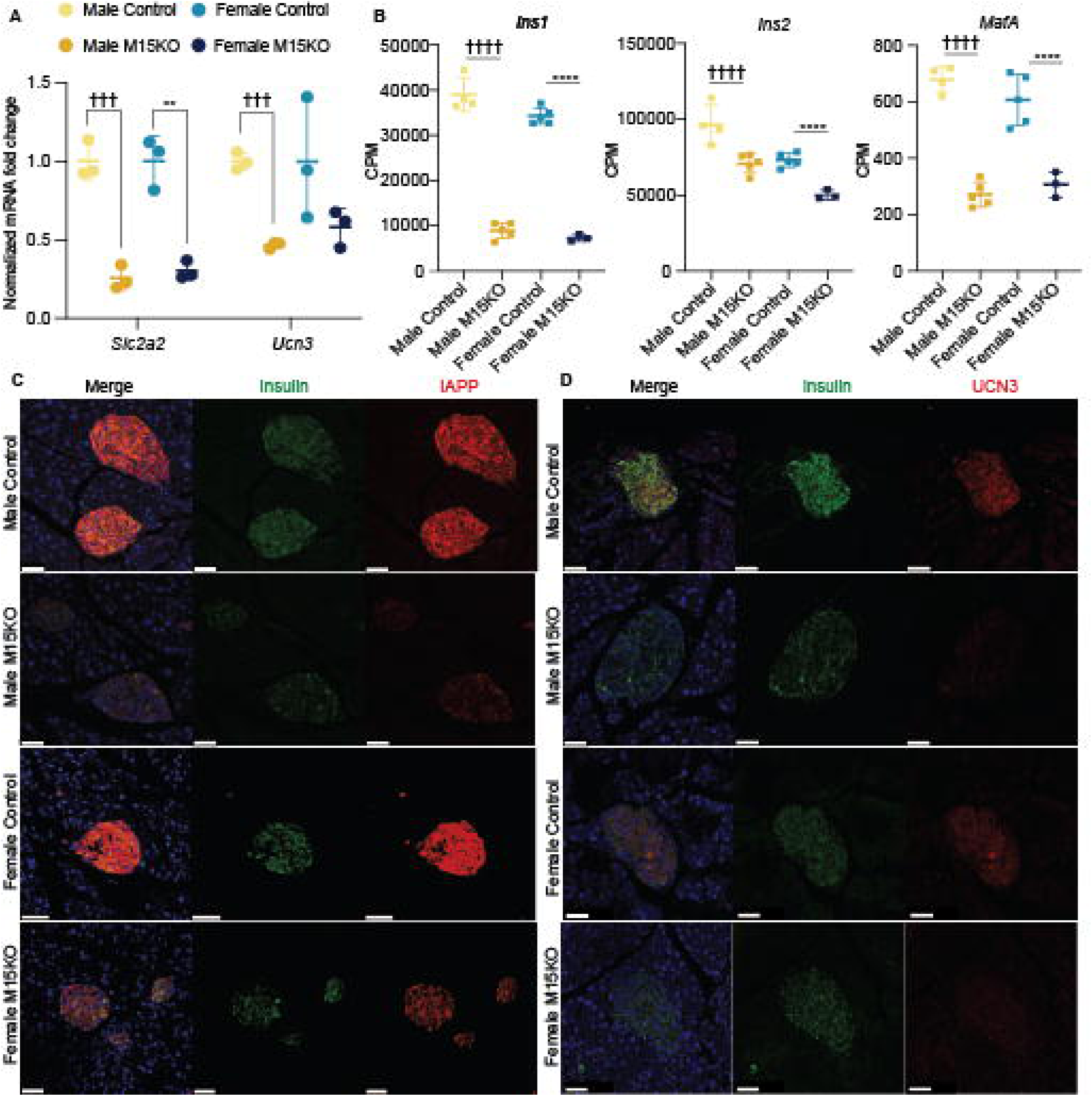
Validation of loss of β-cell maturity gene expression. (A) Graphs of the fold change in expression as determined by RT-qPCR of maturation markers following *Med15* deletion (n=3). (B) Counts per million (CPM) plots of select genes from RNA-seq analysis. (C) immunofluorescence staining of IAPP (red) and (D) UCN3 (red), with insulin (green) and DAPI (blue). Representative images from an n=3 mice are shown. Scale bars are 50um. Data in (A) are mean ± SD and compared using unpaired Student’s t-test. Significance in (B) are p values from RNA-seq analysis. * indicates significance in female mice, † indicates significance in male mice.

### Identification of candidate MED15 interacting transcription factors

Because MED15 regulates gene expression through transcription factors, we hypothesized that deletion of its binding partners would result in similar transcriptional changes as *Med15* knockout. To identify transcription factors that might maintain β-cell maturity with MED15, we compared our RNA-seq dataset to other β-cell-specific, adult-inducible knockout models with whole islet RNA-seq datasets (6,28–31). We compared datasets in two ways. First, we analyzed the overlap of genes regulated by each factor, which revealed that M15KO expression patterns share significant overlaps with all datasets except *Rfx6*KO (Table 1, Supplementary Fig. 7). Among these, the greatest proportional overlap was with NKX6-1 (Table 1), a known binding partner of MED15 (22). PAX6, which is critical for development and maintenance of β-cell identity, had the second highest proportional overlap. This suggests that PAX6 may cooperate with MED15 to maintain β-cell maturity. Across all 7 examined datasets, only *Ucn3*, *G6pc2*, and *Tnfrsf9* were downregulated in all KO models (p<0.05, FDR<1.1, logFC<-0.1). Second, we performed Pearson’s correlation on the datasets of dysregulated genes from each model. The strongest correlations of *Med15* were to Pax6, Rnf20, and Ssbp3 (Fig. 5). Finally, we compared the list of genes regulated by MED15 from our RNA-seq to genes with promoters bound by MED15 or MED1 as identified in a ChIP-seq dataset generated in MIN6 cells (22). Multiple β-cell maturity markers and functional genes that we observed to be downregulated by MED15 loss are bound by MED1 and MED15 (Supplementary Fig. 7). Overall, this shows that MED15 directly regulates key genes involved in β-cell maturation. In sum, our M15KO model shares significant transcriptomic similarities to other models of adult β-cell loss of maturation, and these transcription factors represent candidates for interacting partners for MED15.

**Figure 5:**
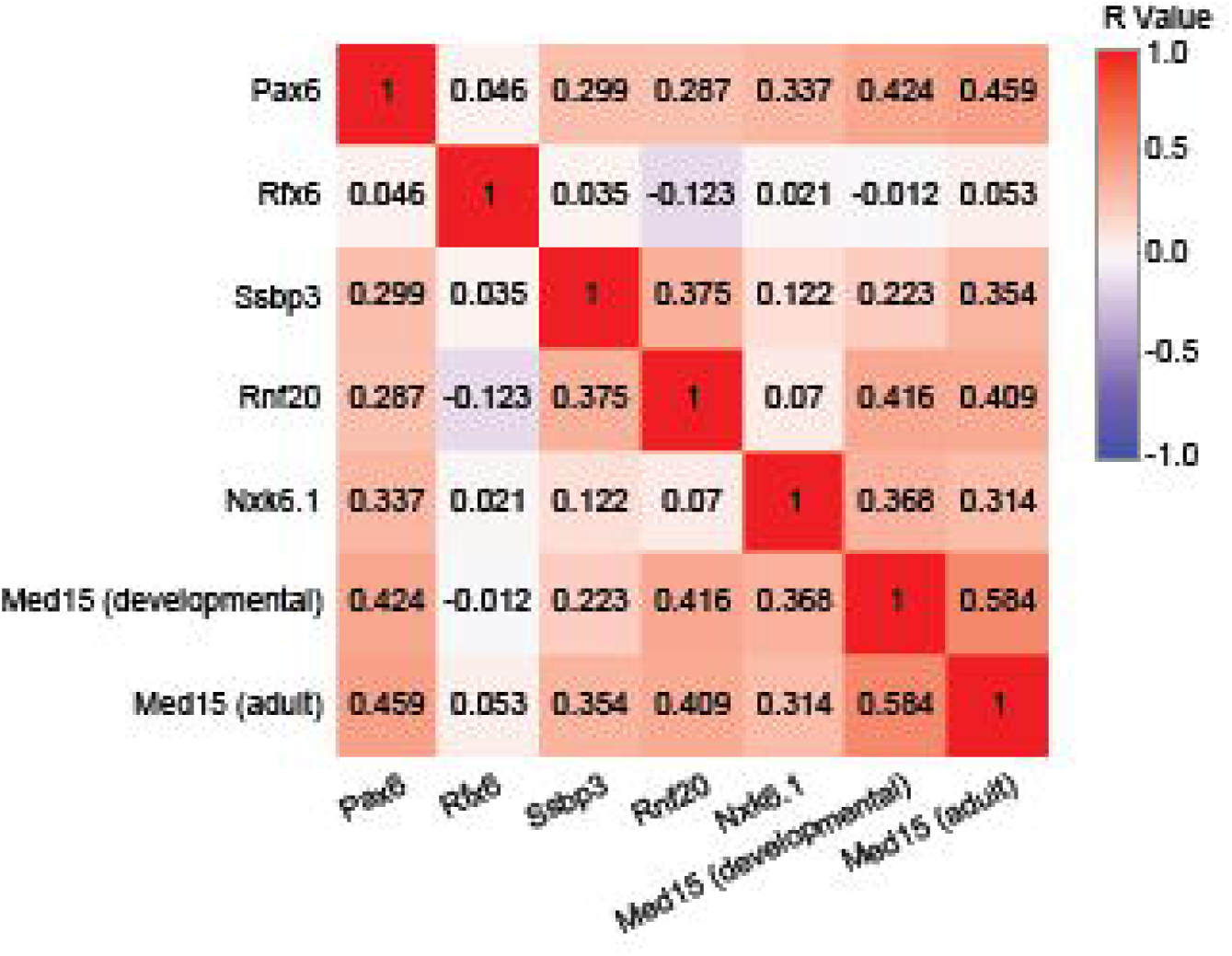
*Med15* KO transcriptomes resemble transcriptomes of other β-cell transcription factor knockout models. Heatmap of Pearson’s correlation between male M15KO RNA-seq dataset and other adult inducible β-cell specific KO models from (6,22,28–31). R values are displayed on heatmap.

**Table 1:**
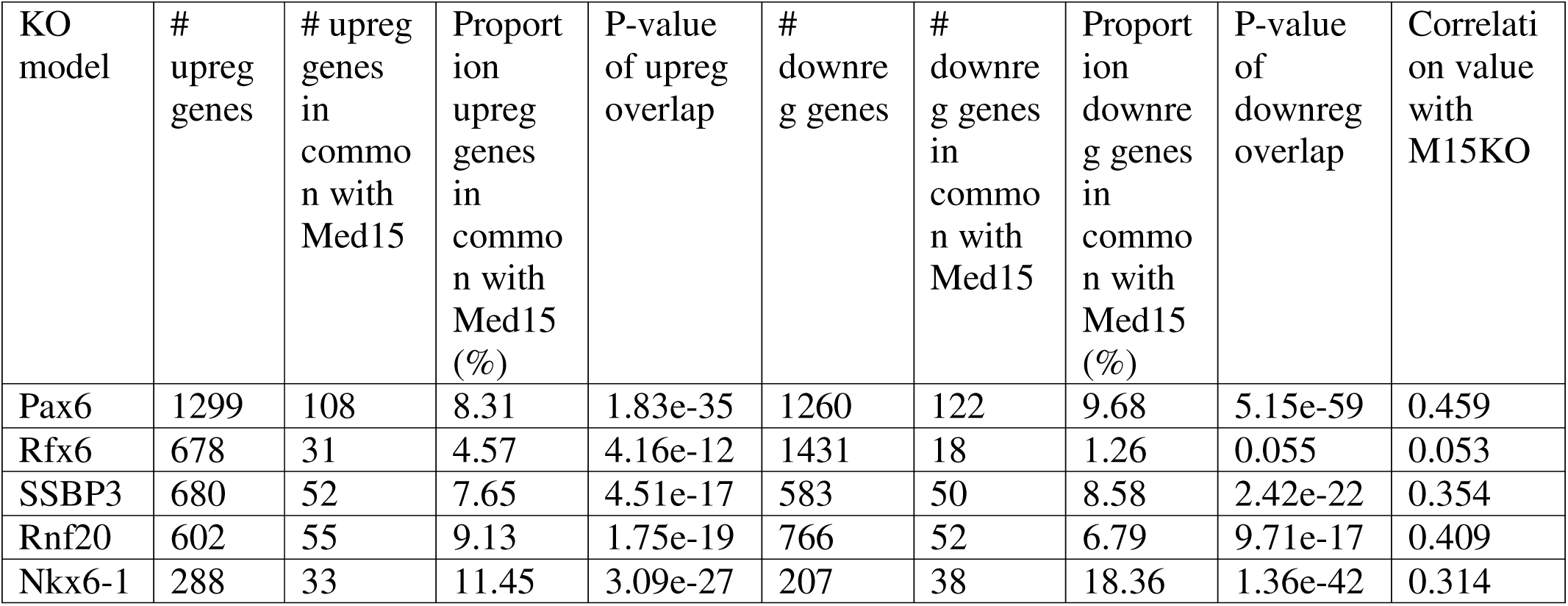
Comparison of genes dysregulated in adult, β-cell specific KO models to genes dysregulated in male M15KO. Columns of up and downregulated gene overlaps corresponds to Venn diagrams in Supplementary Fig. 7. P-value calculated by hypergeometric test. Correlation value from Pearson’s correlation in Figure 5.

| KO model | # upreg genes | # upreg genes in common with Med15 | Proportion upreg genes in common with Med15 (%) | P-value of upreg overlap | # downreg genes | # downreg genes in common with Med15 | Proportion downreg genes in common with Med15 (%) | P-value of downreg overlap | Correlation value with M15KO |
| --- | --- | --- | --- | --- | --- | --- | --- | --- | --- |
| Pax6 | 1299 | 108 | 8.31 | 1.83e-35 | 1260 | 122 | 9.68 | 5.15e-59 | 0.459 |
| Rfx6 | 678 | 31 | 4.57 | 4.16e-12 | 1431 | 18 | 1.26 | 0.055 | 0.053 |
| SSBP3 | 680 | 52 | 7.65 | 4.51e-17 | 583 | 50 | 8.58 | 2.42e-22 | 0.354 |
| Rnf20 | 602 | 55 | 9.13 | 1.75e-19 | 766 | 52 | 6.79 | 9.71e-17 | 0.409 |
| Nkx6-1 | 288 | 33 | 11.45 | 3.09e-27 | 207 | 38 | 18.36 | 1.36e-42 | 0.314 |

### Reducing glycemia does not rescue transcriptional changes in M15KO islets

The expression of some genes involved in β-cell function and insulin secretion can respond to changing glucose levels, especially hyperglycemia. *Txnip* is regulated by glucose concentration, and was highly upregulated in M15KO mice (Fig. 3B-C). To assess if *Txnip* upregulation is influenced by high glucose levels, we incubated isolated control and M15KO islets in high and low glucose media for 4 and 6 hours. After incubation in 5.5 mM glucose, *Txnip* mRNA was expressed at similar levels in islets of both genotypes (Fig. 6A), suggesting that its expression is predominantly regulated by glucose levels.

**Figure 6:**
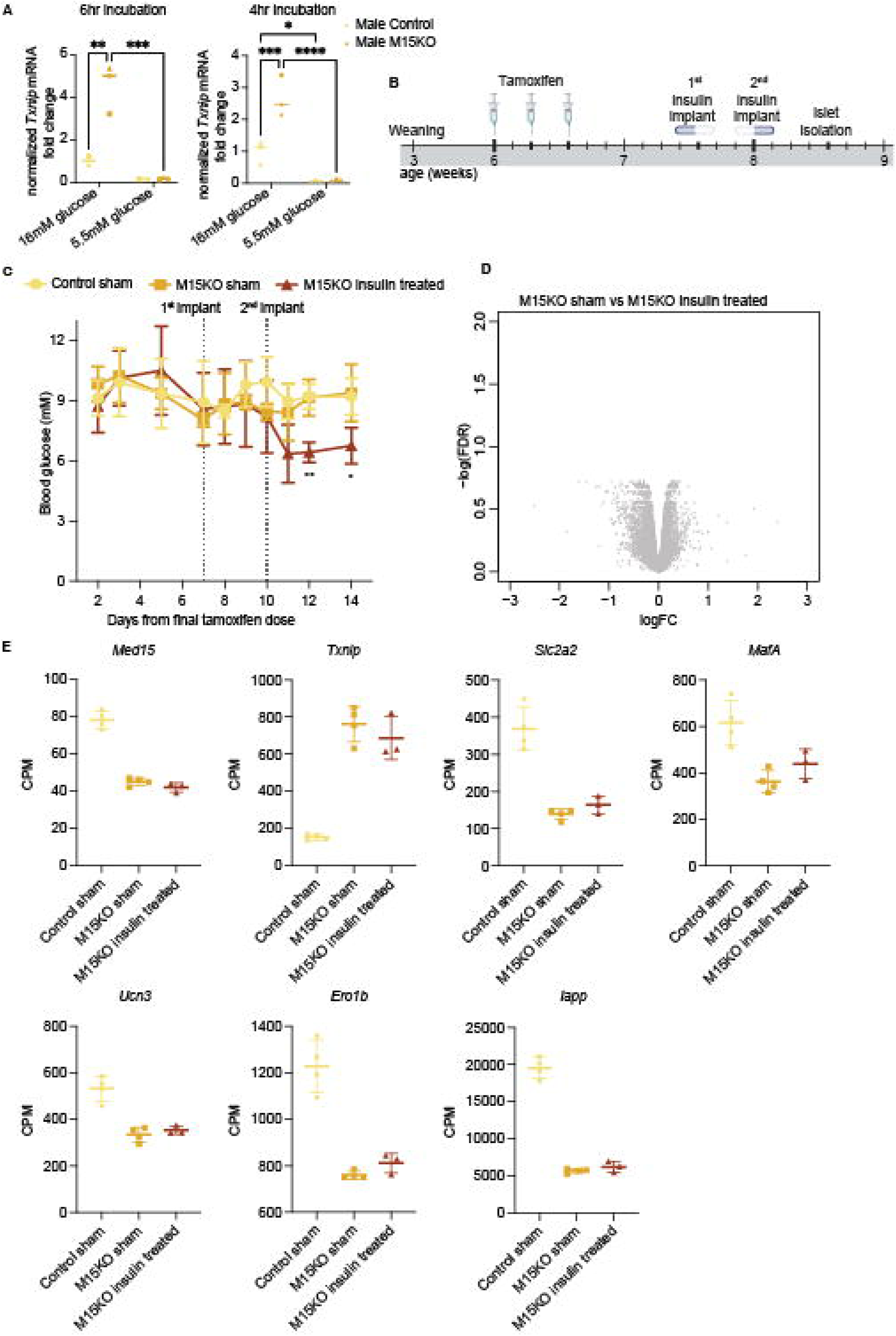
Lowering hyperglycemia does not recover expression of maturity markers in Med15 KO islets. (A) *Txnip* mRNA expression from qPCR of isolated male M15KO and control islets after 4- and 6-hour incubation in high and low glucose media (4hr control, 4hr M15KO, 6hr M15KO n=3; 6hr control n=2). (B) Schematic of timeline for insulin pellet implant M15KO mice. (C) Random fed blood glucose of control untreated, M15KO untreated, and M15KO insulin treated mice. Insulin pellets were implanted on the 7^th^ day after the final tamoxifen dosing, and again on the 10^th^ day. P values indicate two-way ANOVA between M15KO sham and M15KO insulin treated. (D) Volcano plot of DEGs from whole islet RNA-seq of M15KO sham and M15KO insulin treated mice. (E) CPM plots from RNA-seq of whole islet. All genes in KO sham vs KO insulin treated are FDR>0.05. Control sham n=4, M15KO sham n=4, M15KO insulin treated n=3. Data in (A) and (C) are mean ± SD and compared using ANOVA.

Because M15KO mice show impaired glucose control (Fig. 1), we wished to delineate more broadly what transcriptional changes are caused directly by *Med15* loss rather than elevated blood glucose and gradual islet dysfunction. To delineate these regulatory differences *in vivo*, we implanted insulin pellets into M15KO mice seven days after the final tamoxifen dose to lower glycemic levels (Fig. 6B). We initially implanted half an insulin pellet to minimize the risk of hypoglycemia. As blood sugar levels did not decrease after the first implant, another half pellet was implanted three days later. After four days of lowered blood glucose levels (Fig. 6C), we isolated islets from these mice for bulk RNA-seq. Intriguingly, no genes were significantly different between untreated M15KO and insulin-treated M15KO groups (Fig. 6D, Supplementary Table 8). Notably, the major maturation markers *Iapp*, *Slc2a2*, *Mafa*, and *Ucn3*, which are downregulated by *Med15* ablation, remained unchanged in animals receiving insulin treatment (Fig. 6E). This demonstrates that the transcriptional changes described above (Fig. 3) are a direct consequence of *Med15* loss rather than the developing hyperglycemia.

## Discussion

Transcription factors and coregulators play critical roles in pancreas development and function. We previously identified the Mediator subunit MED15 as critical for establishing functionally mature β-cells in mice, but it was unclear if MED15 played a role in β-cells after initial maturation. In the present study, we provide comprehensive evidence that MED15 remains essential in adult β-cells. Specifically, adult mice with β-cell-specific *Med15* loss rapidly develop severe glucose intolerance, elevated fasting glucose, and impaired insulin secretion. RNA-seq revealed downregulation of many β-cell maturation factors and genes involved in insulin secretion. Glucose sensing, metabolism, and insulin folding/processing are all impaired at the transcriptional level. Notably, restoring normoglycemia with pharmacological intervention does not restore maturity at the cellular level, pinpointing MED15 as a key driver of this process. Together with our previous work, this demonstrates that MED15 is critical at multiple junctures in β-cell development and maintenance.

### MED15 regulates genes required for β-cell function in adult mice

We previously showed that MED15 is required for post-natal β-cell maturation (22). Here, we show that MED15’s role in the β-cell does not end at weaning but that it is important throughout life. β-cells must maintain functional maturity to control glucose homeostasis throughout life, and if lost, diabetes can develop (37). Our current study shows that MED15 is required to maintain expression of β-cell maturity and identity markers, including *MafA*, *Nkx6-1*, *G6pc2*, and *Pdx1*. *Med15* is also required to maintain expression of genes required for insulin production and GSIS, such as *Slc2a2*, *Ero1b*, *Cox6a2*, *Sytl4*, and others. Additionally, although M15KO β-cells do not significantly express α-cell markers, many such genes are mildly upregulated. We speculate that this expression may increase over time, potentially reflecting loss of cellular identity and dedifferentiation.

### MED15 cooperates with β-cell transcription factors to regulate β-cell maturity

As a coregulator, MED15 interacts with transcription factors to regulate gene expression. Transcriptional profiling of islets from M15KO mice show that multiple pathways in β-cell function and insulin secretion are impaired after ablation of *Med15*. MED15 likely interacts with transcription factors in the β-cell to control these processes. The phenotype and transcriptional profile of the M15KO mice closely resemble knockout models of certain β-cell transcription factors, suggest that MED15 may bind these transcription factors. When comparing different β-cell KO datasets, we noticed that *G6pc2*, *Ucn3*, and *Tnfrsf9* were the only genes downregulated in all models. *Ucn3* and *G6pc2* are well-characterized β-cell maturation markers, and a *G6pc2* enhancer is bound by islet transcription factors MAFB, NKX2.2, NKX6-1, FOXA2, and HNF4α (38). *Tnfrsf9* (aka Cd137 or 4-1BB) is a costimulatory molecule expressed on immune cells that inhibits cancer progression via P38MAPK/PAX6 signaling. In non-obese diabetic mice, *Tnfrsf9* KO suppresses diabetes development by reducing the pathogenicity of β-cell reactive T cells (39), but why this TNF receptor is strongly regulated by so many β-cell transcription factors and co-regulators is unclear and warrants further study. *Tnfrsf9* may be involved in maturation or represent a new marker of β-cell maturity.

RNA-seq also showed that both insulin genes were downregulated in the M15KO islets, although to different extents. *Ins1* expression was reduced five-fold, whereas *Ins2* expression was downregulated only 1.6-fold (Fig. 3E). This is similar to differential insulin expression seen in *Neurod1* β-cell KO mice (5), and notably, MED15 physically interacts with NEUROD1 (22). Thus, MED15 may regulate *Ins1* by binding NEUROD1.

### MED15’s effect on the β-cell transcriptome is direct and not glycemia-mediated

*Txnip* is among the most highly induced genes in M15KO islets (Fig. 3B-C), particularly in males, which exhibit more severe glucose intolerance. *Txnip* is strongly induced by high glucose (40), and we hypothesized that this may reflect an indirect response to hyperglycemia rather than a change due to *Med15* loss. Indeed, some phenotypes of M15KO mice might be caused by hyperglycemia. We tested this hypothesis by implanting insulin pellets into M15KO mice to lower their blood glucose levels, but, after four days of lowered blood glucose, M15KO mice receiving insulin implants had no transcriptional changes compared to M15KO mice with sham treatment. Because improved glycemia did not rescue the maturation defects or other transcriptional changes in M15KO mice, these changes are likely directly due to *Med15* loss. This supports a model whereby MED15 continuously interacts with β-cell transcription factors to maintain maturation and function.

### Insulin production and processing is impaired in M15KO mice

Genes strongly downregulated in M15KO mice include those involved in insulin secretion, e.g. *Erol1b*, which is required for folding of proinsulin (41), calcium activated cation channel *Trpm5* (42), *Sytl4*, which encodes granuphilin and is required for insulin granule docking (43), melanophilin (*Mlph*), which functions in insulin granule exocytosis (44), and *C2cd4a*, which regulates glycolytic genes (45). *Atp2a2*, which encodes for SERCA2, was also strongly downregulated in M15KO islets. SERCA2 maintains endoplasmic reticulum Ca2+ concentrations, and its deletion in mice causes glucose intolerance due to impaired maturation of insulin processing enzymes (46). Taken together with the impaired insulin secretion in response to KCl (Fig. 1I-J), this suggests that defects in insulin production at both a transcriptional and protein processing level may contribute to the impaired glucose control in M15KO mice.

### MED15 maintains β-cell maturity in both sexes

Diabetes prevalence, glucose homeostasis, and β-cell function are sex dependent in humans and rodent models. We observed glucose intolerance in both male and female M15KO mice, with more severe presentation in males. Our M15KO mice are on a C57/B6J background, which has sexual dimorphism in glucose homeostasis, namely improved glucose tolerance in females (33,47). Thus, the differences seen in the M15KO male and female mice are likely attributable to innate sex differences of this strain. Importantly, both sexes exhibited similar transcriptional changes, especially of loss of β-cell maturity and function; therefore, MED15 likely regulates and maintains β-cell maturation in both males and females.

### Therapeutic significance

We show that MED15 is required to maintain β-cell maturation after weaning. β-cell maturation is of interest in the context of stem-cell derived β-cells that may be used for transplantation into people with type 1 diabetes. Although current *in vitro* protocols can differentiate insulin-producing β-cells, their function and transcriptome are dissimilar from mature, primary β-cells (48). Additionally, β-cell dedifferentiation and loss of maturity are mechanisms of β-cell failure that can lead to diabetes (37). Therefore, better understanding the networks that not only initiate, but also maintain β-cell maturation, is critical to furthering diabetes therapeutics.

## Acknowledgments

We thank Derek Dai for help with mouse surgeries, Cuilan Nian for help with islet isolations, and Mitsuhiro Komba for assistance with islet perifusion. We are grateful to Paul Sawchenko for providing rabbit anti Ucn3 antibody. Imaging work was supported by the Imaging Core (RRID:SCR_026573) at BC Children’s Hospital Research Institute (BCCHR).

## Author Contributions

Conceptualization, FCL, CBV, and ST; Methodology, RJS, SPC, FCL, and ST; Investigation, RJS, MD, and SPC; Writing – Original Draft, RS; Writing – Review & Editing, RS, SPC, MD, FCL, CBV, and ST; Funding Acquisition, CBV, FCL and ST; Supervision, FCL and ST. ST takes full responsibility for the work as a whole, including study design, access to data, and the decision to submit and publish the manuscript.

## Prior Presentation

Parts of this study were published in abstract form at the European Association for the Study of Diabetes Annual Meeting in Vienna, Austria, September 2025. This manuscript has been deposited in bioRxiv.

