## Supplemental Material for "Transcriptional coactivator MED15 is required to maintain β-cell maturity"

**Online Supplemental Material.**

**Supplementary Methods**

**Mouse genotyping primers**

Mice were genotyped with the following primers: *Med15* forward: TAAGGTGCTGTGTGTGGTTGGC, reverse: GTAATGATGAGGCTAAAAGGGCTG; *mTmG* wildtype forward: CTCTGCTGCCTCCTGGCTTCT, wildtype reverse: CGAGGCGGATCACAAGCAATA, mutant reverse: TCAATGGGCGGGGGTCGTT; *Pdx1^CreER^* forward: CGCAAGAACCTGATGGACATG, reverse: GCTACACCAGAGACGGAAATC.

**RNA isolation and RT-qPCR**

Islets were placed in RLT buffer (79216; Qiagen) containing 1% β-mercaptoethanol (M6250; Sigma-Aldrich) and RNA extracted according to manufacturer protocol with a Qiagen RNeasy Mini Kit (74104; Qiagen). cDNA was made using ThermoFisher High-Capacity cDNA Reverse Transcription kit (4368814; ThermoFisher Scientific) and 25ng of cDNA template was used per reaction with Taqman Fast Advanced Master Mix (4444557; ThermoFisher Scientific). The following pre-designed Taqman Gene Expression Assays were used: *Slc2a2*, Mm00446229_m1; *Ucn3*, Mm00453206_s1; *Actb*, Mm02619580_g1; *Gusb*, Mm01197698_m1.

*Med15* and *Txnip* qPCR was performed as described (22) using the following primers and probes tagged with FAM, ZEN, and Iowa Black FQ (IDT Technologies). *Med15* (Forward: GCATGGCTGTGGTGTCTA; Reverse: CCTGCTGTTGCTGGAATTG; Probe: CAACAGCAGCAGCAGCAACAACAA). *Txnip* (Forward: AAGGATGACTTTCTTGGAGCC; Reverse: ACATTATCTCAGGGACTTGCG; Probe: TTTGAGGATGTTGCAGCCCAGGA).

qPCR was performed on a QuantStudio 5 (Applied Biosystems) and analyzed using the Threshold (Ct) method.

**RNA-sequencing and analysis**

Sample quality control was performed using an Agilent 2100 Bioanalyzer system. Qualifying samples (RNA integrity number >8) were prepared following the standard protocol for the llumina Stranded mRNA prep (Illumina). Sequencing was performed on an Illumina NextSeq2000 with Paired End 59bp×59bp reads at the Sequencing Facility of the UBC School of Biomedical Engineering (https://bme.ubc.ca/home/sequencing-facility/). We sequenced >20 million reads per sample. The raw FASTQ files from the facility were concatenated and trimmed using Trimmomatic version 0.36 (1) with parameters LEADING:3 TRAILING:3 SLIDINGWINDOW:4:15 MINLEN:36. Next, trimmed reads were aligned to the Ensembl reference genome GRCm39 (GCA_000001635.9) (<https://may2025.archive.ensembl.org/Mus_musculus/Info/Index>) using Salmon v1.41 (2) with parameters -l A -p 8 --gcBias. Transcript-level read counts were imported into R and summed into gene-level read counts using tximport (3). Genes not expressed at a level >1 count per million (CPM) reads in at least three samples were excluded from further analysis. The gene-level read counts were normalized using the trimmed mean of M‐values (TMM) in edgeR (4) to adjust samples for differences in library size. Differential expression analysis was performed using the quasi-likelihood F-test with the generalized linear model (GLM) approach in edgeR (4).

**Immunostaining**

Tissue was blocked in 5% horse serum (26050-088; Gibco) and incubated overnight at 4°C with primary antibodies, and for two hours at room temperature with secondary antibodies (Supplemental material). Coverslips were mounted with Vectashield PLUS (H-2000; Vector Laboratories). Sections were imaged using a confocal microscope (Leica SP8; Leica Microsystems).

For β-cell area quantification, slides were treated with Vector® TrueVIEW® Autofluorescence Quenching Kit (SP-8400-15) following secondary antibody incubation, imaged on a BX61 widefield fluorescence microscope, and tiled using the cellSens Dimension software (Olympus). CellProfiler v4.2.5 was used to quantify images (33). Briefly, after thresholding each channel, cell nuclei were differentiated with watershedding, and glucagon-positive (GCG+) and insulin-positive (INS+) cells were counted and normalized to total pancreatic nuclei.

We used the following antibodies:

| **Target** | **Catalogue Number; Vendor** | **Dilution** |
| --- | --- | --- |
| Insulin | IR002; Dako | 1:4 |
| Glucagon | G2654; Sigma | 1:1000 |
| MED15 | 11566-1-AP; Proteintech | 1:50 |
| UCN3 | gift from Sawchenko lab | 1:400 |
| IAPP | 4157; BMA Biomedicals | 1:150 |
| FITC anti-mouse | **715-095-150;** Jackson Immunoresearch | 1:200 |
| FITC anti-rabbit | **711-095-152** | 1:200 |
| Alexa Fluor 647 anti-guinea pig | 706-605-148), | 1:200 |
| Cy3 anti-rabbit | 711-165-152 | 1:200 |
| DAPI | 4083S; Cell Signalling Technology | 1:2000 |

**Supplementary Tables**

**Supplementary Table 1.** Individual p-values for all figures.

**Supplementary Table 2.** Differentially expressed genes from RNA-seq of male M15KO and control islets.

**Supplementary Table 3.** Differentially expressed genes from RNA-seq of female M15KO and control islets.

**Supplementary Table 4.** Genes regulated in both post-natal *Ins1-Cre* *Med15*KO and adult *Pdx1-CreER* M15KO islets (p<0.05, logFC<-0.5 or >0.5).

**Supplementary Table 5.** Gene set enrichment analysis (GSEA) analysis from M15KO islet RNA-seq. The Kyoto Encyclopedia of Genes and Genomes (KEGG), WikiPathways (WP), Biological Processes (BP), and Reactome Pathway (RA) databases were used.

**Supplementary Table 6.** GSEA of pathways involved in endoplasmic reticulum stress and unfolded protein response.

**Supplementary Table 7.** Differential expression of genes involved in the ER-UPR.

**Supplementary Table 8.** Differentially expressed genes from RNA-seq of whole islets from untreated versus insulin treated M15KO islets.

**Supplementary Table 9.** Genes bound and regulated by MED15 and genes bound by MED1 and regulated by MED15.

**Supplementary figures.**

**Supplementary Figure 1. *Med15* mRNA and protein are downregulated after KO.**

(A) CPM plot from RNA-seq (male control n=4, male M15KO n=5, female control n=5, female M15KO n=3) and (B) RT-qPCR gene fold changes (n=4) showing loss of *Med15* expression in whole islets of M15KO mice. (C) IF staining from M15KO and control islets show loss of MED15 in M15KO islets. Data are mean ± SD. Significance in (A) are p values from RNA-seq analysis. Data in (B) are compared using unpaired Student’s t-test.

**Supplementary Figure 2. Islet architecture is unchanged after Med15 KO.** (A) quantification of GCG expressing cells in the islet core (defined as more than 2 cells from exterior). Male control, male M15KO, female control n=3, female M15KO n=2.

**Supplementary Figure 3.** **Male and female M15KO islets share dysregulated genes. (A) Multidimensional scaling plots of islet RNA-seq samples from M15KO and control mice of each sex. (B)** Venn diagram show shared differentially expressed genes (FDR < 0.05, |logFC| ≥ 0.5) between male (yellow) and female (blue) M15KO islets.

**Supplementary Figure 4. Mediator subunit expression is not affected following *Med15* deletion.** RNA-seq CPM plots showing relative expression of Mediator complex subunits from the (A) head, (B) tail, (C) middle, and (D) kinase modules in M15KO and control (CTRL) islets.

**Supplementary Figure 5. Integrated stress response is not activated in M15KO islets.**

Heatmap of M15KO and control islets gene expression from RNA-seq showing expression of integrated stress response (ISR) genes.

**Supplementary Figure 6.** **β-cell immaturity genes are upregulated in M15KO islets.**

Heatmap showing M15KO and control islet RNA-seq expression of genes associated with β-cell immaturity. This “immature” cluster of genes downregulated during β-cell maturation was generated from single-cell RNA-sequencing analysis in (37).

**Supplementary Figure 7.** **MED15 regulates similar genes as other β-cell transcription factors.** Venn diagrams of overlapping up and downregulated genes between M15KO and other adult β-cell transcription factor KO models (6,28-31).

**Supplementary Figure 8.** **MED15 binds and regulates β-cell function and maturation factors.** Euler diagrams of genes both bound by MED15 and regulated in M15KO versus genes bound by MED1 and regulated by MED15. Upregulated genes are in blue and downregulated genes in orange circles. Selected downregulated and bound genes in both males and females are highlighted. Full list of
